# On the permeability of cell membranes subjected to lipid oxidation

**DOI:** 10.1101/2020.11.30.403345

**Authors:** Daniel Wiczew, Natalia Szulc, Mounir Tarek

## Abstract

The formation of transient hydrophilic pores in their membranes is a well-recognized mechanism of permeabilization of cells exposed to high-intensity electric pulses. However, the formation of such pores alone is not able to explain all aspects of the so-called electroporation phenomenon. In particular, the reasons for the sustained permeability of cell membranes, which persist long after the pulses’ application, remain elusive. The complete resealing of the cell membranes takes indeed orders of magnitude longer than the time of electropore closure as reported from molecular modelling investigations. A possible alternative mechanism to explain the observed long-lived permeability of cell membranes, lipid peroxidation, has been previously suggested but the theoretical investigations of membrane lesions, containing excess amounts of hydroperoxides, have shown that the conductivities of such lesions were not high enough to reasonably explain the entire range of experimental measurements. Here, we expand on these studies and investigate the permeability of cell membrane lesions that underwent secondary oxidation. Molecular dynamics simulations and free energy calculations on lipid bilayers in different states show that such lesions provide a better model for post-pulsed permeable and conductive electropermeabilized cells. These results are further discussed in context of sonoporation and ferroptosis, respectively a procedure and a phenomena, among others, in which alike electroporation substantial lipid oxidation might be triggered.

**Highlights:**

1. The contribution of secondary lipids’ oxidation to the permeabilization of model membranes is quantitatively assessed
2. Small patches of secondary lipids’ oxidation cause formation long-lived pores in lipid bilayers.
3. The cholesterol content of membranes enhances the life-time of the formed pores.
4. A single pore accounts for the measured post-pulse electropermeabilization of cells.
5. The diffusion of the secondary oxidation lipids, even after pores closure leads to permeability of lipid membrane.

## Introduction

Electroporation (EP) is a well-known bio-method used to enhance the permeability of biological cell membranes [1–4]. This technique, which consists of exposing cells to pulsed electric fields (PEFs), enables the transport of various molecules such as drugs and genes, across their lipid membrane [5–8]. A current goal in improving our understanding of electroporation is the development of a comprehensive microscopic description of the phenomenon. Information about the sequence of events describing electroporation is gathered from measurements of electrical currents through planar lipid bilayers along with the characterization of molecular transport of molecules into (or out of) cells subjected to electric field pulses and from atomistic level details provided by Molecular Dynamics (MD) simulations of model membranes. It is now well accepted that long and intense electrical pulses induce rearrangements of the membrane components (water and lipids), which ultimately lead to the formation of aqueous hydrophilic pores whose presence increases substantially the ionic and molecular transport through the otherwise impermeable lipid bilayers. One of the main unsolved questions related to the electroporation phenomena is that the induced permeability of cell membranes can persist up to minutes after the applied electric fields are terminated as reported by several studies [9–12]. This is in large discrepancy with the pores sustainability that MD simulations indicate, which are rather in the order from tens of ns to the μs time scale [13,14].

It has been suggested over two decades ago, that membranes can be oxidized when subject to conditions similar to that of electroporation-based technologies. There is experimental evidence indeed that pulsed electric fields can increase the extent at which unsaturated lipid acyl chain peroxidation occurs. In particular, it has been shown that the application of external electric fields alters the phospholipid composition and properties of liposomes, vesicles, and cells [15–21]. The presence of oxidized lipids within bio-membranes is known to modify their physical properties and, in particular, their permeability [22–25]. One cannot therefore exclude that molecular uptake following PEFs treatments may, at least partially, take place through diffusion across oxidized/permeabilized lipid bilayer patches and not solely across the so-called electropores. In cells, the repair of such patches is a long-time process that might explain the sustainability of membranes’ leakage.

Several studies demonstrated that applying PEFs to cells induces generation of reactive oxygen species (ROS) and oxidative damage of unsaturated lipids, as confirmed for instance by increased concentration of conjugated dienes [4] and hydrogen peroxide [19,20]. ROS concentrations and the extent of lipid peroxidation increase with electric field intensity, pulse duration, and the number of pulses (see [4] for review). PEFs don’t generate radical oxygen species by direct action on the media exposed but appear to enhance their generation [26–28], most probably following a destabilization of mitochondria membrane or Fenton reaction [29,30].

Among the most potent ROSs, the hydroxyl radical (HO•), the superoxide radical anion (O_2_·), and the hydroperoxyl radical (HOO•) are short-lived and highly reactive and, therefore, are believed to play a prominent role in cell membranes’ lipid peroxidation [31]. These radicals can initiate chain reactions leading to chemical oxidation of the lipids and thus change in the physical properties of cell membranes [32,33].

HO· and HOO· radicals if reaching the interior of membranes containing unsaturated lipids (**L**) can initiate primary oxidized products by an allylic hydrogen abstraction (see Fig. 1) leading to the formation of lipid radical **L**· [33,34]. **L**· reacts with molecular oxygen, which is highly abundant in the membrane’s interior [31] to form lipid peroxide radicals **LOO**·. The latter may further abstract hydrogen from another lipid to form a primary peroxidation product, **LOOH**, (hydroperoxide), and a lipid radical **L**·, initiating thereof a chain reaction.

**Fig. 1.**
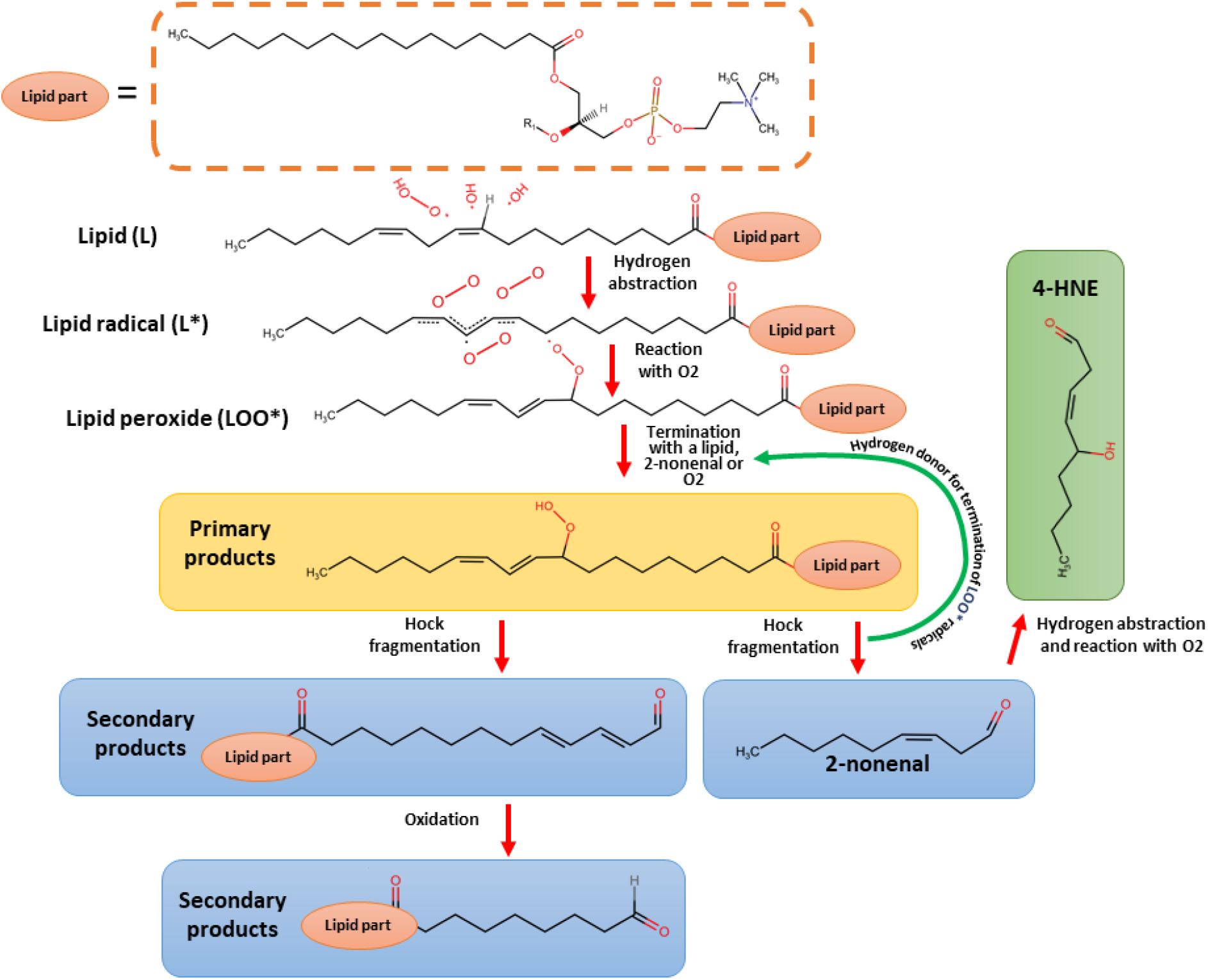
Schematic pathway of process formation primary and secondary oxidation products.

In a recent work [35] we have investigated the permeability of membranes containing patches of various concentrations of hydroperoxidized lipids mimicking therefore the presence of patches of lipids that underwent peroxidized chain reactions. In particular, we estimated their conductance and permeability to monovalent ions using MD simulations and free energy calculations. The results of the calculations were compared to experimental measurements on electropermeabilized cells. Our data showed that the permeability and conductance increase dramatically by several orders of magnitude in hydroperoxidized bilayers However, this increase was not sufficient to reasonably account for the entire range of experimental measurements. Furthermore, our data, consistent with experiments on giant unilamellar vesicles (GUVs) containing up to 60% lipid hydroperoxides or exclusively lipid hydroperoxide species [36], showed that the bilayers preserve their membrane integrity.

In this paper, we extend our investigation to study the permeability of membranes that underwent secondary oxidation. Indeed, oxidative lipid damage can result in various products with truncated lipid tails ending with either an aldehyde or carboxylic group [37,38]. Hydroperoxides are converted into secondary products like 2-nonenal and PoxnoPC by Hock fragmentation [39]. The 2-nonenal is further converted to 4-HNE (4-hydroxynonenal) by reacting with a radical and oxygen in a similar fashion as unsaturated bonds in unsaturated lipids, thus potentially contributing to the kinetic’s oxidation process (positive feedback loop). Previous MD simulations showed that oxidized lipids with an aldehyde group disturb the bilayer more than the ones with a peroxide one [40]. Bilayers with aldehyde-truncated tails on the other hand undergo spontaneous pore formation within a few hundred ns and lead in some cases to the bilayer complete disintegration (micellation) [40],[41],[42]. In contrast, Runas and Malmstadt [33] reported formation of pore defects in GUVs containing only 12.5% aldehyde-truncated 1-palmitoyl-2-(9¢-oxo-nonanoyl)-sn-glycero-3-phosphocholine (PoxnoPC). Spontaneous pore formation in GUVs was reported also by Sankhagowit *et al.* [43] under conditions where aldehyde-truncated lipids were produced.

Our aim here is to thoroughly investigate the properties of lipid membranes that underwent such drastic chemical changes, with regard to their permeability. We will specifically consider a model lipid bilayer reflecting properties of a real system (highest fraction of lipids is POPC) [44] with a central patch of secondary product lipid oxidation (see Fig. 2).

**Fig. 2:**
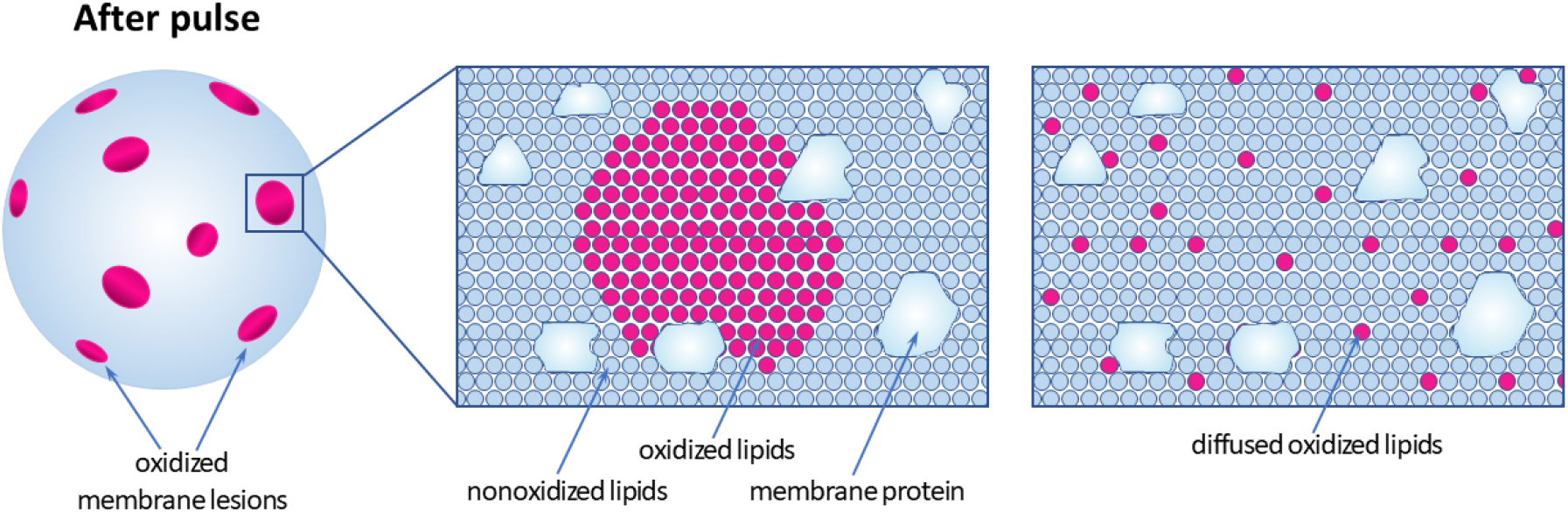
Schematic representation of oxidized membrane lesions (secondary oxidized lipids), which are expected to be formed in the cell membrane after exposure to electric pulses. The schematic is hypothetical and the lesions are not drawn to scale. The image in the center schematically depicts the molecular organization in one of the lesions. The one on the right represents the state after a certain time, where the secondary oxidized lipids diffused out (adapted from [35]).

The presence of such patches in cell membranes results from the radical chain peroxidation reaction that takes place between neighboring lipids. Fast kinetic reaction ratios obtained from state-of-the-art quantum chemistry calculations suggest that degradation of primary to the secondary oxidation products might be faster than lateral diffusion of lipids [39]. To mimic crowded conditions in the cellular environment, and the scenario where the patches are created in a gel-like raft domain of the cell wall, we considered as well, a second model membrane of POPC with 40% cholesterol content. Both models (with and without cholesterol) contain a central patch of PoxnoPC, a stable byproduct of PLPC [45] or POPC oxidation [46]. These simulations allowed to directly follow for microseconds the features of such patches. Furthermore, similarly to our previous investigation [35], we used free energy calculations applying the Unified Free Energy Dynamics (UFED) method [47] to quantify the conductance and permeability to simple monovalent ions of bilayers in which oxidized lipids have diffused out of the initial patch to form homogeneous domains (see Fig. 2 right). We show how such analyses is used to compare the properties of these model systems to experiments conducted on cells.

## 2. Materials and methods

### 2.1 Patch system calculations

#### 2.1.1 Systems

The initial system (SYST I) was built using PACKMOL [48] and consisted of a large 1-palmitoyl-2-oleoyl-sn-glycero-3-phosphocholine (POPC) bilayer, in which the central patch was replaced by a 1-Hexadecanoyl-2-(9-oxo-nonanoyl)-sn-glycero-3-phosphocholine (PoxnoPC) bilayer of 64 molecules per leaflet (see S2). PoxnoPC is a zwitterionic oxidized phospholipid bearing an aldehyde function at the end of its truncated sn-2 acyl chain. The whole bilayer model was further solvated in a solution containing sodium, calcium, and chloride ions (for composition details see S2 in the SI). The second system (SYST II) was the analogous previous one, but the POPC lipid bilayer contained a 40 mol % of cholesterol (see S2 in the SI)

Additionally, four simple lipid bilayer systems (POPC) containing PoxnoPC at molar concentrations of 0%, 10%, 20%, and 50% (respectively BIL1 to BIL4) were set and embedded in NaCl solutions (see Table S3.1 in the SI for more details). A System with the concentration of 80% of oxidized lipids was not stable (data are not shown here). BIL1 to BIL4 were studied to investigate how local concentrations of oxidized lipids after their diffusion out of the central patch (Fig. 2 right) influence the permeability and conductance of the lipid membrane. The systems were built in a similar way to SYSTI and SYSTII using PACKMOL, except that PoxnoPC lipids were distributed randomly across the POPC lipid bilayer. Over 1μs nanoseconds of simulations were run for every concentration to extract the corresponding features and properties.

#### 2.1.2 MD Simulations

To perform the simulations, POPC and cholesterol were parameterized with the CHARMM36 lipid force field [49]. We developed a CHARMM-consistent Force field parameter for modeling PoxnoPC following the protocol used by Jeffery B. Klauda et al. using state-of-the-art quantum chemistry calculations [49] (see S1 in the SI for the detailed protocol). We used the TIP3 model [50] for water and the electronic continuum correction (ECC correction) for the ions representation [51] as it allows an accurate description of sodium and calcium ions interaction with lipid bilayer’s lipid headgroups (see [35]).

All simulations were performed using the GROMACS software considering 3-d periodic boundary conditions. All systems were first minimized using the steepest descent algorithm to remove interatomic clashes, then equilibrated at constant temperature T (310 K), and constant (semi-isotropic) pressure P (1 atm). Bonds between heavy atoms and hydrogens were constrained using the LINCS algorithm [52] allowing the use of a 2 fs MD timestep. The long-range electrostatic interactions were evaluated using the Particle Mesh Ewald method [53]. The Fourier grid spacing for the Particle Mesh Ewald method was optimized at the beginning of each simulation, to get the highest performance during the simulation. A switching function was used between 0.8 and 1.2 nm to smoothly bring the short-range electrostatic interactions and the van der Waals interactions to 0 at 1.2 nm.

#### 2.1.3 MD analyses

To track the changes in the lipid bilayer’s BIL1 to BIL4 properties during the simulations, the thickness and area per lipid (APL) was assessed using the PyBILT software [54].

To compare the influence of cholesterol on the lipid bilayer fluidity, the lateral diffusion D coefficient of the PoxnoPC lipids was estimated using the Stokes-Einstein’s equation that links D to the mean square displacement of molecules. Hence, in SYST I and SYST II, ten lipids were selected based on the criteria of being not involved in the oxidized lipid clusters (see below). Then the diffusion coefficient was calculated using the GROMACS’ MSD tool for diffused lipids for the last 40 ns of simulation time. The sizes of the pores reported were roughly estimated by visual inspection using the grid projected onto the membrane with Visual Molecular Dynamics (VMD) [55] “grid” tool in the ruler extension.

### 2.3 Free energy calculations using the Unified Free Energy Dynamics

Free energy methods allow ones to calculate free energy profiles with respect to a collective variable (CV). The CV maps a high dimensional atomic coordinate of an explored process (conformational change, ion permeation) into a low dimensional representation, that can be manipulated to obtain desired results. In the study, we used the Unified Free Energy Dynamics [56],[47] approach. The latter combines ideas from two methods, d-AFED (driven Adiabatic Free Energy Molecular Dynamics) and metadynamics, resulting in superior convergence in comparison to both methods. UEFD was successfully used in our previous works [35],[57],[58] to obtain the potential of mean force (PMF) profile of ions and small radical species along the normal to a lipid bilayer. d-AFED works by connecting a CV *q* with harmonic potential (with coupling constant **κ)** to an extended variable s (see equation (1)). The extended variable is adiabatically decoupled from the rest of the system, which is achieved by choosing large **κ** and mass of the extended variable. Such treatment allows the extended variable *s* to be set to a temperature T_s_ higher than that of the system to ensure better sampling of states in the phase space. The temperature T_s_ is usually set such that k_b_T (k_b_ is Boltzmann constant) is similar to the energetic barrier of the studied process. Thus, the extended variable can easily cross the barrier and drags the CV alongside. In UFED, the regions of the phase space are explored using a bias that ensures a better sampling of the infrequently explored phase space regions. The bias, similar to the one used in metadynamics is in the form of Gaussian hills with given height h and variance sigma, that is deposited gradually along the extended variable trajectory. The PMF Φ(*s*) profile can be recovered by numerically integrating from the force *F(s)* acting on the extended variable *s*.

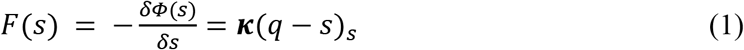

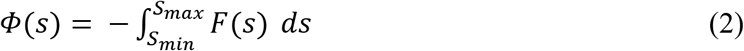

where *s* and *q* are extended variable and collective variable respectively and ***κ*** denotes the coupling constant for the mentioned harmonic potential. In our study, the extended variable was defined as the center of the lipid bilayer, whereas the collective variable was defined as the position of an ion (sodium or chloride) with respect to the bilayer center.

#### 2.3.1 System preparation for UFED

BIL1 to BIL4 were equilibrated in the semi-isotropic NPT ensemble (T= 310 K; P= 1 atm) for over a μs. The bilayer thickness and area per lipid were calculated to verify convergence of the equilibration, calculated [59]. The electrostatic potential profile along the membrane normal (z-axis) was evaluated considering the last 100 ns of the simulation using the 1D Poisson’s equation,

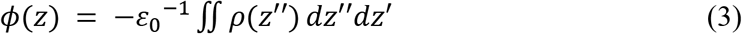

Where *ρ(z)* is the molecular charge distribution along *z*. The integration was performed by slicing the system into 1000 bins and averaging charge density within each bin and then integrating it according to the above equation using the GROMACS *g-potential* tool.

#### 2.3.2 UFED parameters

One of our work’s interests was obtaining the PMF profile of the translocation of an ion (sodium and chloride) along the direction normal to the lipid bilayer containing secondary oxidation product lipids. Therefore, the CV was defined as a z-position of the ion. In agreement with our previous investigation, this collective variable was coupled with a harmonic constant of 10^4^ kJ/mol/nm to a meta-variable of mass m_s_ = 2.10^4^ a.m.u. The extended system was coupled to a bath at a temperature T_s_ = 400 K using a generalized Gaussian moment thermostat (GGMT). Gaussian-shaped potential hills, with a height of 1.0kJ/mol and width of 0.01 nm, were deposited every 5000 MD steps (100 fs). To prevent an overall movement of the lipid bilayer in the z-direction, the center of mass of the system (estimated from the average z position of the Lipids head group phosphorus atoms) was restrained in the z-position using a harmonic potential with a spring constant of 3500 kJ.mol^−1^nm^−2^.

#### 2.3.3 UFED simulation protocol

The free energy calculations were performed using GROMACS 4.6.3 [60] patched with PLUMED 1.3 [61] that includes the implementation of d-AFED/UFED free-energy calculations [56],[47]. The parameters for the molecular dynamics were similar to the system preparation phase (2.3.1 above), except for the LINCS algorithm [52] parameters, which were adjusted toward more accurate calculations (“lincs-order” set to 6 and “lincs-iter” set to 2). The temperature of the simulation was set T = 300 K both for steering and UFED calculations (see paragraph below), and thermostat coupling constants were set to 1.6 ps. More details about these parameters can be found in the supplementary material S3.4.

A multiple walkers’ strategy, which involves the parallel evolution of multiple (sixteen) metadynamics simulations sharing the same bias potential history file, was employed in order to increase computational efficiency. The UFED calculations were hence performed using 16 separate sub-simulations (called further walkers) in parallel. Each walker started at a different initial configuration, in which the position of the ion (Na or Cl) along the bilayer normal was obtained by steering the latter toward the center of the lipid bilayer in an independent MD run. The initial configuration of this run was the last frame from the well equilibrated (1 μs) trajectory described in section 2.3.1. The steering was performed using a spring constant of 3500 kJ.mol^−1^nm^−2^ and a velocity of 0.3 nm/ns using PLUMED. The steered configurations were saved by every 0.1 nm, from 6.5 nm to 3.45 nm to the center of the lipid bilayer, giving 16 configurations in total. Further molecular dynamics parameters for steering were the same as for UFED, the same goes for temperature that was set to T = 300 K.

The UFED calculations were performed for 80 ns for each walker of the systems listed in the 2.3.1 (8 calculations in total - 4 for chloride, 4 for sodium). The output for the UFED variables (q, s, etc&) were saver every 0.04 ps of the simulations. An example of two trajectories for two walkers is shown in the SI S3.4.1. To ensure convergence and estimate uncertainty, a second UFED run was performed in the same way as the first one. Except that, the initial configuration was different than in the first run.

#### 2.3.4 Analysis of PMF profiles

The PMF profiles were evaluated based on the equation (3) using a custom Matlab scripts (Matlab R2019a, MathWorks) which were adapted from the scripts developed by M. A. Cuendet [35],[47],[57]. This was achieved by a multistep calculation workflow that started by dividing obtained force along a collective variable *s* into 120 bins in the interval z = [-3.0, 0.0] nm. Then the forces *F* = *κ(q* − s)_s_ were sampled over the UFED run and averaged within each bin. These forces then were slightly smoothed with a kernel smoothing regression (Matlab function ksr) with a bandwidth of 1/30 nm. Finally, the forces were integrated with cumulative trapezoidal numerical integration (Matlab function cumtrapz) to obtain the PMF over the interval z = [-3.0, 0.0] nm. This profile was then mirrored across the center of the bilayer.

To estimate the uncertainty of the profiles, we performed an additional UFED run for each of the investigated systems. The second UFED run was performed in the same way as the first run, except that the initial configurations of the walkers were different. The PMF profiles were determined the same way as the previous run. A mean over the two runs is shown as a PMF in the manuscript (see Fig. 6). The two runs are used to confirm the convergence and reliability of the UFED method see S3.5.2 in the SI.

### 2.4 Estimations of the permeability and the conductance

The bilayer permeability to a given ion was calculated according to the inhomogeneous solubility-diffusion model as in (4) [62–64].

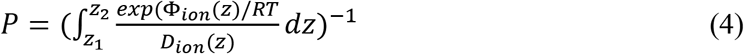

where Φ*_ion_(z)* is the PMF (Na or Cl) ion along the z-axis, *R* is the universal gas constant, *T* the temperature ow the system and *D_ion_(z)* the diffusion coefficient of the ion along the z-axis (see next section). The integration boundaries were taken as *z*∈[-3.0,0.0] *nm.* The uncertainty of the values was estimated given the uncertainty of the PMF and that on the *D_ion_(z)*. The former was estimated based on the maximum and minimum value of obtained permeation from two separate UFED runs. The lower and upper two values were multiplied by 1.33 and 0.67, corresponding to the maximum variation between maximum/minimum values relative to smoothed *D_ion_(z)* profile (more in the SI 4.1). The uncertainty is about +/− 33% in the average scenario (highest error among all diffusion calculations). It was applied to all other diffusion calculations.

As in [35], the total and ionic conductance (in S/m^2^) of Na and Cl were calculated using [62],[65]:

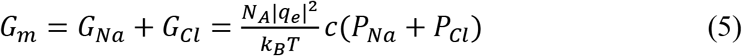

where *G_m_ G_Na_* and *G_cl_* are respectively the total sodium, and chloride conductance. *N_A_* is Avogadro’s number, *q_e_* is ion’s charge, *k_B_* and *T* are Boltzmann’s constant and temperature. The concentration and temperature for the sake of this work were assumed as 150 mM and T = 300 K respectively – the same as used in the UFED and diffusion calculations. The uncertainties were calculated based on the uncertainty propagation rule for addition, diffusion’s uncertainty, and two UFED calculations, obtaining lower and upper bound for conductivity in a similar manner as in the case of permeation.

### 2.5 Diffusion coefficients

The diffusion coefficients of the ions across the membranes were determined similarly as in [35],[62] by performing umbrella sampling on the ion constrained in the z-positions using a harmonic potential with a spring constant *κ* = 3500kJ.mol^−1^nm^−2^. The initial configuration was obtained from the both steered trajectories performed during preparation to the UFED calculations (see 2.3.3 above), thus giving 32 configurations in total. The MD parameters were the same as in the UFED calculations (see section 2.3.1). For each walker *i* ∈ [0, 31], a 10 ns long trajectory was generated, where the first two ns were considered as an equilibration phase. The remaining 8 ns were divided into four equal segments. For each one, the diffusion coefficient for a given configuration *i* was calculated as:

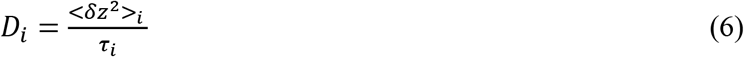

The variable *τ_i_* here is the correlation time for the i-th configuration and was calculated using the method of Hummer [66]. Further, *D_i_* obtained for each of the configurations was averaged over the four 2 ns long parts. Finally, the averaged values were smoothed with a kernel smoothing regression (Matlab function ksr) with a bandwidth of 0.2 nm. The smoothed profiles were used in the calculation of membrane permeability and conductivity.

## 3. Results

### 3.1 Patches of secondary products of lipids oxidation undergo spontaneous and long-lived pore formation

Fig. 3 reports snapshots from the MD trajectories for SYSTI and SYSTII (respectively Pure POPC and POPC:40% cholesterol). The systems represent small patches of PoxnoPC located in the center of the membrane that model “hot spots” resulting from a peroxidation chain reaction initiated by a ROS attack, followed by a secondary oxidation of the hydroperoxide lipids (see Fig. 1).

**Fig. 3.**
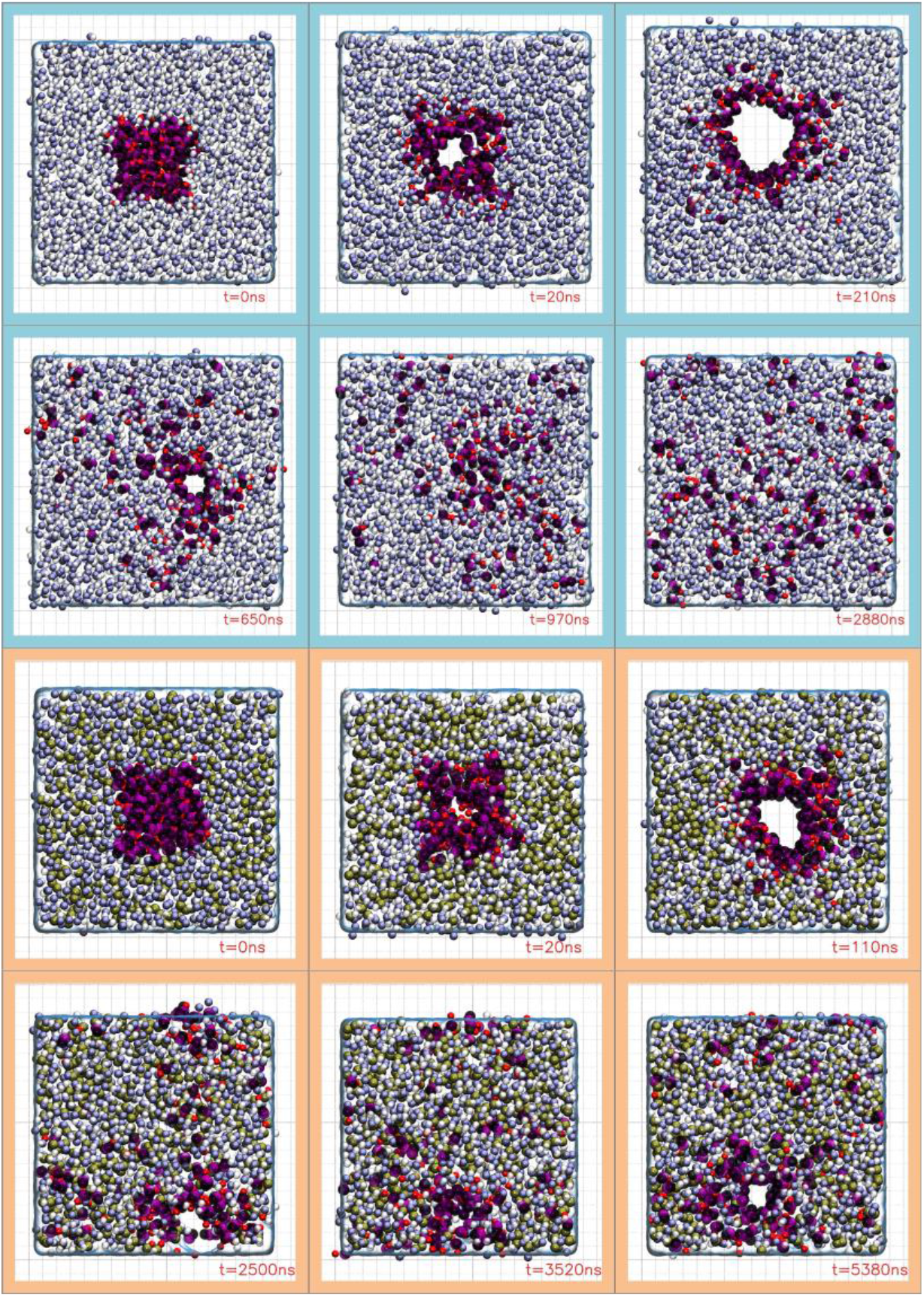
Time evolution of pore formation in the bilayer systems (top view) without cholesterol (blue panels) and with cholesterol (orange panels). The grid size is 1nm. The blue and white spheres represent the choline and phosphate groups of the POPC molecules’ zwitterionic head groups. The magenta and black spheres represent the choline and phosphates groups the PoxnoPCs molecules’ head groups, the red spheres correspond to aldehyde oxygen of the PoxnoPCs lipid tails, and the cholesterol’s OH groups are drawn as yellow spheres. The corresponding membrane cross sections can be found in SI S2.3 along with the videos of the full MD trajectories.

The simulations indicate that the presence of such a PoxnoPC patch, given the localized high concentration of aldehydes, leads within a few nanoseconds to a pore formation across the bilayer. As the simulations are carried out at 0 surface tension (anisotropic NPT ensemble at 1 atm, see Casciola & Tarek [7]), these pores expand quickly to reach a maximum size of about 7 nm and 5 nm in diameter for SYST I and SYST II respectively. Such pores are therefore large enough to enables transport of DNA [67],[68], and fluorescent dyes such as: Propidium Iodide, or Calcein. For SYST I, because of the liquid crystalline state of the POPC, the PoxnoPC molecules diffuse out of the patch during the ~3 μs MD simulations, leading to a pore shrinking, basically as the number of PoxnoPC molecules forming the pore decreased. Within a couple of μs, this pore shrinks hence to a diameter of ~ 0.6 nm, then the pore closes. For SYST II, the diffusion of the PoxnoPC out of the initial patch was much less rapid, due to the gel-like dynamics of the remaining lipid bilayer (40% cholesterol). Indeed, estimation of the lateral diffusion coefficients D of PoxnoPC in the system with 40% cholesterol (6.2±0.9) × 10^−12^ m^2^s^−1^, was smaller than that of the pure POPC bilayer, namely (12.0 ± 2.8)× 10^−12^ m^2^s^−1^. Consequently, the number of oxidized lipids around the pore remains high enough to maintain an aperture of ~ 1.35 nm diameter 5 μs down the trajectory.

### 3.2 Properties of lipid bilayer after diffusion of the PoxnoPC lipids

In order to study the permeability of the regions of cell membranes that contain oxidized lesion long after the initial state, we have modeled several systems for which the secondary products have diffused enough so that the pores close (see for instance last panel Fig. 3 of SYST I). These systems model larger regions of cell membranes composed therefore of a homogeneous distribution of PoxnoPC within an intact bilayer. We set hence POPC bilayers with concentrations of PoxnoPC ranging from 10 mole% to 50 mole% and studied both their characteristics and permeability.

First, MD runs exceeding 100 ns for every system studied did not reveal any destabilization of the membranes at these oxidized lipids concentrations. No pores/water or ion leaks were formed during the simulations. The structural properties of the bilayers reported in Fig. 4 show monotonic changes of the area per lipid and of the bilayer thickness compared to the values for the pure POPC bilayer (respectively 3.93 nm and 0.65 nm^2^). The latter are within 1% of the experimental values reported for POPC at 30C [69] which provides some confidence in the force field parameters used.

**Fig. 4.**
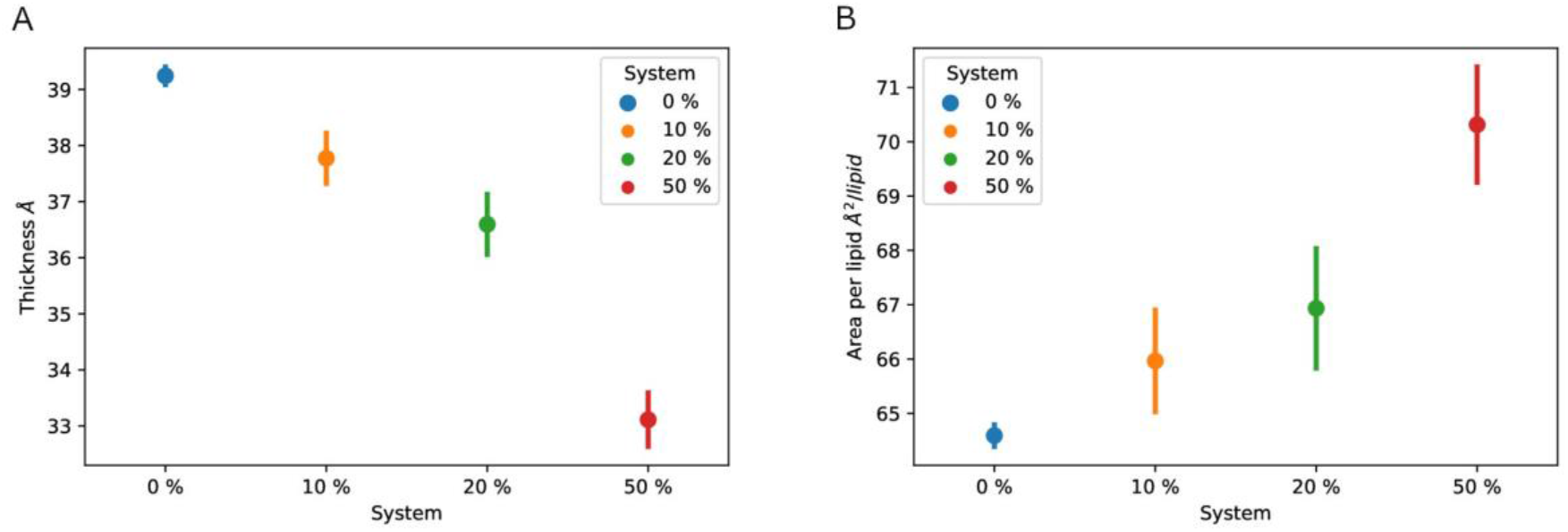
Thickness (A) and average area per lipid (B) for lipid membranes with 0%, 10%, 20% and 50% percent of secondary oxidation lipids (PoxnoPC) in the POPC lipid bilayer. The bars indicate standard deviations based on the last 100 ns of the MD trajectories.

### 3.3 Ion penetration barrier across the bilayer after the diffusion of the PoxnoPC lipids

The PMF profiles of the translocation of Na^+^ and Cl^−^ ions across the bilayer systems with 0% (pure POPC bilayer) 10%, 20%, and 50% homogeneous distribution of PoxnoPC are reported in Fig. 5. All the profiles are Λ-shaped, similar to those reported in previous studies [35],[70]. As the percentage of the oxidized lipids increases, the free energy barrier progressively decreases for both the anion and the cation, as reported for lipids bilayers containing hydroperoxides [35]. As expected, the bilayer with 50% PoxnoPC lipids presents the lowest barrier to diffusion in comparison to the other bilayer systems.

**Fig. 5.**
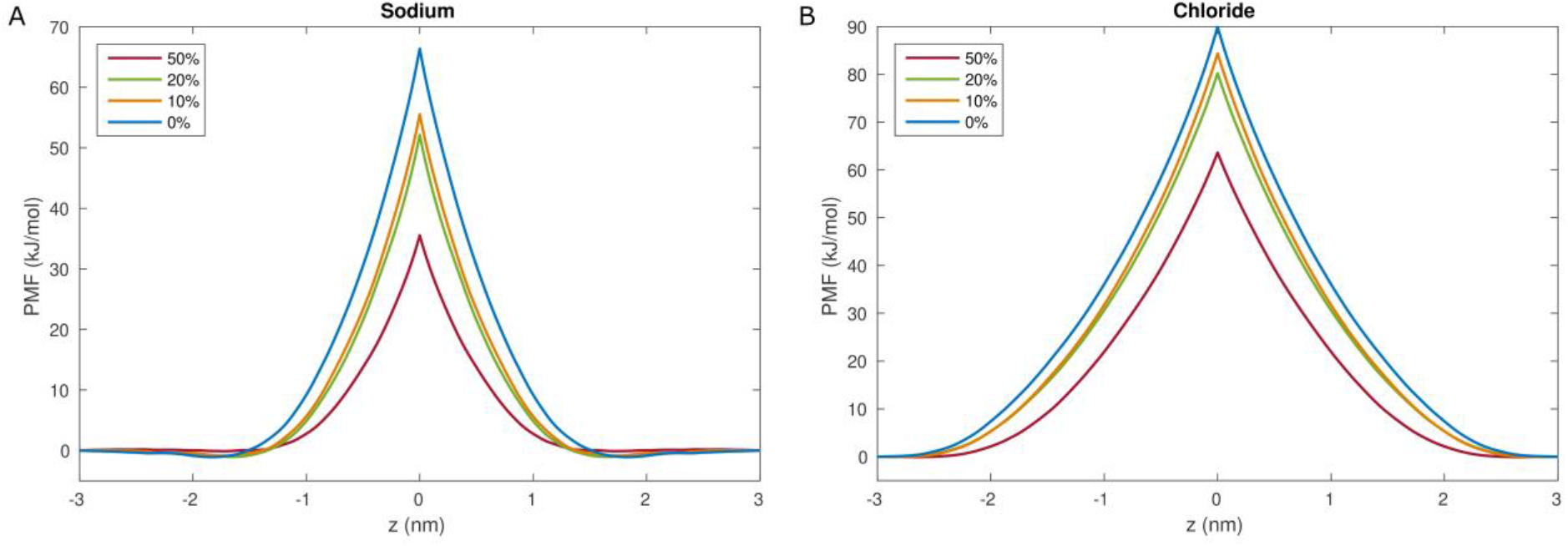
The PMF profiles of Na^+^ ion (A) and Cl^−^ ion (B) in the investigated bilayer systems, BILL1 to BILL4, namely POPC with percentages (drawn in different colors) of homogeneous distribution of PoxnoPC. The center of the bilayer is located at z = 0 nm.

Here, the calculations with concentrations higher than 50% of oxidized lipids were not considered due to formation of pores in the simulations of such systems (for instance at 80% PoxnoPC), as was the case for SYST I and SYST II. It is fair to assume accordingly that systems with oxidized lipid concentration higher than 50% would have negligible or virtually zero free energy barrier to permeation as ions can pass freely through the formed pores.

### 3.4 Permeability and ion conductance of lipid bilayer with secondary oxidation products

The permeability to each ion species and the total conductance of the systems BILI to BIL4 are depicted in Fig. 6. As the percentage of peroxidized (PoxnoPC) lipids increases, the permeability and conductance do so as well. Noticeably, the Na^+^ permeability of the POPC lipid bilayer with 50% PoxnoPC is 5 orders of magnitude higher than the permeability of the pure POPC bilayer, increasing from 10^−11^ m/s to 3.10^−6^ m/s. The same can be observed for the Cl^−^, where the permeability increases from 2.10^−15^ m/s to 6.10^−11^ m/s, *i.e.* by over 4 orders of magnitude.

**Fig. 6.**
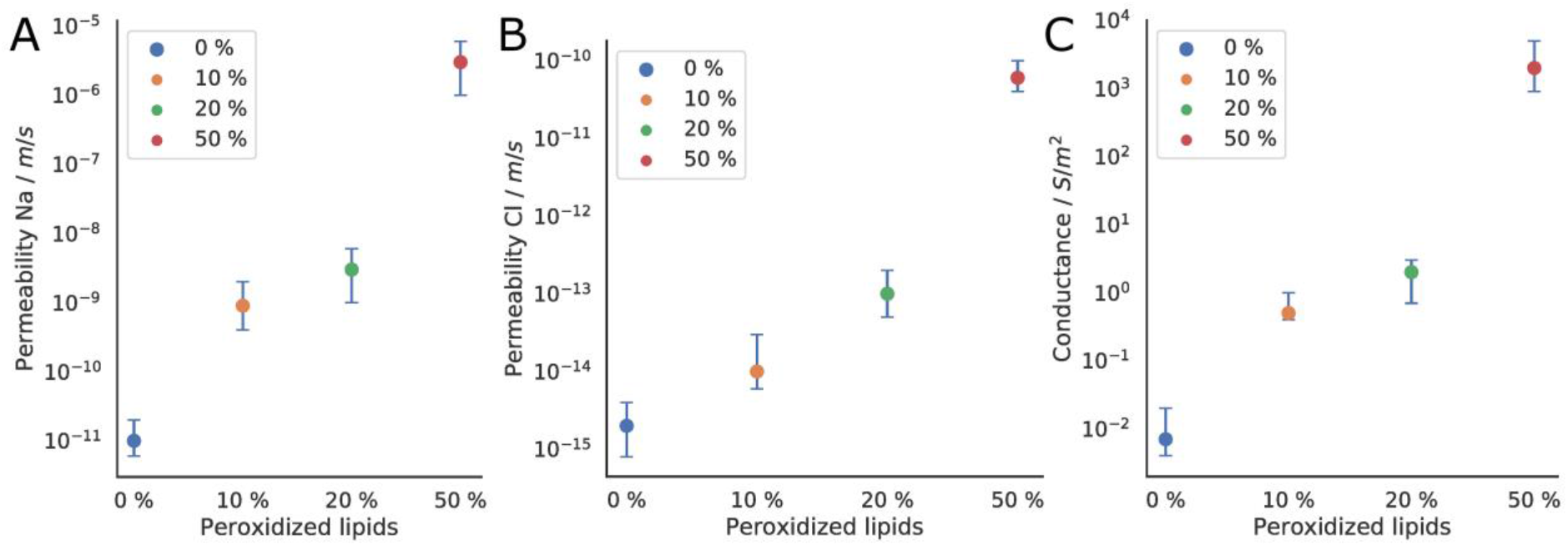
Permeability to Na^+^ (A) and Cl^−^ (B) ions and overall conductance (C) of the BIL1 to BILL4 studied systems.

These changes are reflected in the magnitude of changes in the overall conductance. Hence, the analysis of the simulations data using eq (6) shows that the conductance of the lipid bilayer containing 50% of secondary oxidation PoxnoPC lipids product is 5 orders of magnitude higher than that of the non-oxidized lipid bilayer.

## 4. Discussion

It has been suggested in recent literature that lipid oxidation could be a possible reason for sustained cell membranes permeability, a phenomenon that manifests itself as persistence of permeability following treatment with high electric field, and that lasts up to minutes after the electric fields that trigger a so-called membrane electroporation are switched off. As we will discuss later, this phenomenon may not only be encountered in electroporation [71] but also whenever cells or tissues are exposed to high levels of ROS, e.g. subject to ultrasounds triggering sonoporation [57] or to photodynamic therapy [72].

To date, however, no rationale for this phenomena has been offered. Previous studies by our group have concluded that membranes that underwent primary oxidation alone, namely membranes with an excess amount of hydroperoxides, were not leaky enough to account for cell permeability and conductance measured in experiments where cells were exposed to PEF usually used in electroporation-based technologies and treatments. The aim of the present study was to investigate whether oxidative lesions of cell membranes composed of secondary lipid oxidation products can give rise to such a behavior.

Hence, we have herein conducted MD simulations of molecular systems aiming at modeling the behavior of cell membranes subjected to lipid oxidation. The first set of simulations was meant to model membranes immediately after exposure to the electric field and therefore to peroxidation. We have assumed that peroxidation of lipids followed by secondary oxidation would produce “spots” or “patches” of oxidized lipids at the surface of cells (Fig. 2). The presence of such patches in cell membranes results from a radical chain peroxidation reaction that takes place between neighboring lipids, once the process is initiated for instance by the attack by a ROS [73]. We have studied the outcome of such spots when they are formed in a fluid lipid phase, namely a POPC lipid bilayer above its phase transition temperature, or when formed instead in a “gel-like” domain modeled here as a POPC bilayer containing a high amount of cholesterol.

### 4.1 Secondary oxidation lipids and pore formation in the lipid bilayer

The results of Section 3.1 show that a small patch of secondary oxidation aldehyde lipids (PoxnoPC) is enough to form spontaneously a large pore (up to ~ 5 nm diameter) in both lipid bilayers - with and without cholesterol. This indicates that the oxidation of only a small area of a cell membrane would induce the formation of wide enough pores to transport ions and large molecules. Similar behavior was observed in other MD investigations where a high concentration of various oxidation aldehyde lipids induced pore formation and micellization [40],[74]. In an experimental study of giant unilamellar vesicles (GUVs), Runas & Malmstadt reported that 12.5% or higher concentration of PoxnoPC in a GUV induces fluorescein-dextran uptake and suspected that it could be due to pore formation [75]. Sankhagowit *et al.* [43] reported that GUVs with light-induced lipid oxidation show line tensions changes and surface area per lipid changes, that could be interpreted as a pore opening.

### 4.2. Secondary oxidation lipid’s pores and their conductance

While commonly described electric field-induced pores (also called electropores) following rearrangement of the membrane components are easily attributed to the membrane damage allowing ionic and molecular transport by computational as experimental studies [4], [7], [76] they do not explain sustained (long time scale) post-pulse permeation.

Among the experimental methods which enable the detection of lipid oxidation products one may cite spectrophotometric [77,78], chromatographic [79], and immunochemical [80] ones. Other methods using fluorescent dyes are widely applied as well to probe oxidation in live cells and organisms [81,82]. However, these techniques are able to detect only the average content of oxidized species and cannot hence distinguish between uniform and localized distribution of oxidized lipids [83]. The sensitivity of these methods remains a challenge, certainly not enough to detect a few “spots” of secondary oxidation products. In addition, most oxidized lipids stay probably within the membrane and only the low-molecular products, like malondialdehyde that dissociate are detected in many of such detection methods.

In the following we will question first and foremost if pores that are formed by oxidative patches similar to those studied above, can be, in anyway, correlated to the sustained permeability and conductance measured in cells exposed to PEFs. We recall from [35] that *N_pore_* the number of pores needed in a cell to provide an increase of normalized membrane conductivity Δ*g*(in units nS/pF) can be expressed as:

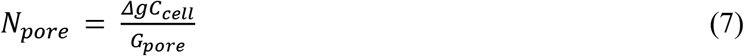

where *C_cell_* is the total capacitance of the cell membrane and *G_pore_* the pore’s conductivity. The analytical expression for cylindrical pore can be expressed [84] as:

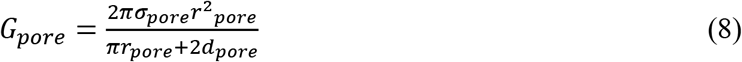

where *σ_pore_* = (*σ_extra_ − σ_intra_)/ln(σ_extra_/σ_intra_*), is a function of σ_*extra*_ and *σ_intra_*. the extra- and intracellular *r_pore_* is a diameter of pore and *d_pore_* its length.

Using equation (7) and (8), one can, therefore, estimate the number of pores required to reach a given change in cell’s conductivity Δ*g* or the change Δ*g* per pore (Δ*g*/*N_pore_*)

Hence, considering the pore radius *r_pore_* ~ 2.5 nm (data from section 3.1) and the values of *C_cell_* ~ 6 *pF* [85], *σ_extra_* = 1.48 *S/m* [86] and *σ_intra_* = 0.5 5/m extracted from experiment, and *d_pore_* ~ 5 nm, the thickness of the bilayer, we find that Δ*g*/*N_pore_* amounts to ~0.3 nS/pF per pore.

Quite interestingly, the conductivity changes reported for GH3 cells from 5 to 10 s after the application of 60 ns PEFs of amplitudes ranging from 4.8 to 14.6 kV/cm is in the range of 0.1 - 1.5 nS/pF (values obtained from Fig 3a. of [87] normalized with the GH3 cell’s capacitance of 6 pF [85]).

Hence, our data indicate clearly that one to few pores of the size of those formed by the small patch of oxidized lipids in the GH3 cells is enough to account for the changes reported in their membrane’s conductivity. The mere lipid patch of 6.6 x 6.6 nm of 64 truncated oxidized PoxnoPC lipids per leaflet mimicking the presence of patches that underwent peroxidized chain reactions is able to form a pore big enough to explain most of the change in ion conductivity, and thus increased permeation.

### 4.3 Pore lifetime in a gel like lipid bilayer models

In the above discussion we highlighted that pores formed by small patches of oxidized lipids could explain the high permeability of cells. However, the timescale during which pores forming in our simple model of fluid-like bilayers (POPC) remain open is too short (few microseconds) to account for all experimental observation. To mimic crowded conditions in the cellular environment, and the scenario where these patches are created in a gel-like raft domain of the cell membrane, we considered enriching lipid domains composed of highly fluid lipids (like POPC) with cholesterol, decreasing thereby their fluidity [88]. We set up a similar lipid bilayer as previously, but with the addition of 40 mol % cholesterol. As stated in Section 3.1, the oxidized lipids of the patch quickly form pores and under these conditions, the lifetime of the pore increased significantly. In our simple model the pore remains open during the whole simulation time (~ 5 μs) investigated here showing that the membrane fluidity and the location of the “oxidative spots” might significantly affects pore’s lifespan.

### 4.4. Conductance after truncated oxidized lipid diffusion and pore closure

The simulations of SYST I and SYST II show that oxidized lipids diffuse out of the pore rim toward the neighboring region where they form domains with high “homogeneous” concentrations of truncated lipids. These concentrations decrease with time (due to further lateral diffusion of PoxnoPC) while the domains enlarge. In order to study the permeability of such regions, we estimated the conductance (see Section 3.4) for model (POPC) membranes with 50%, 20%, 10% PoxnoPC with experimental values of conductance normalized to the GH3 cell’s 6pF capacity. We first calculated the normalized conductance’s of the 3 systems using the capacitance C_m_ per unit area (~ 1 μF/cm^2^ [91]) which resulted in 200 nS/pF, 0.2 nS/pF, 0.05 nS/pF. (for more information see [35]).

Quite interestingly, the modeled conductance increase *Δg_ox_* is comparable to the conductance increase *Δg_exp_* reported for GH6 cells and recorded 2-3 min after subject them to a 60 ns single PEF [87], namely ~0.05 nS/pF to ~0.5 nS/pF with respect to control. We can estimate the fraction *f_ox_* of the cell’s membrane that is needed to obtain the measured conductances (more details and derivation in [35], section 4.1) using:

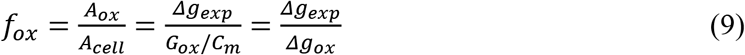

Accordingly, agreement with the data reported for GH6 cells would require that about 0.025% to 0.25% of the cell’s lipids membrane area to be oxidized at 50%, whereas for the other limiting case with 10% degree of oxidation, it would require 100% to 1000% of the lipid membrane area to be oxidized.

Since pore re-forming was observed during the simulation with excess amounts (over 80%) of homogeneous PoxnoPC distribution, it could be hypothesized, that a mixture of pores and regions with a high concentration of oxidized lipids could also lead to significant cell’s sustained permeation.

### 4.5. Lipid oxidation and pore formation beyond electroporation

In the introduction of this paper, we have indicated that evidence suggests that PEFs don’t generate radical oxygen species (ROS) directly, but appear to enhance their intracellular or extracellular generation [26–28]. The scenarios presented above are therefore not restricted to electroporation-based treatments. They might be encountered whenever ROS generation is enhanced, in which case one would expect sustainable pore formation due to secondary lipid oxidation. Most obviously, one should expect similar pore occurrences when cells and tissues are subject to photodynamic therapy. PDT is a therapeutic modality in which photosensitizers excited at specific wavelengths generate reactive oxygen species [90]. As a result, the target cells loaded with such photosensitizers undergo damage that can lead up to necrosis or apoptosis partly due to PDT induced lipid oxidation and its consequences on membrane permeability (see [91] for a recent review).

In the present discussion one would like to focus yet on another biomedical techniques beside electroporation or PDT, known to enhance lipid oxidation, namely sonoporation.

In recent years, research in the field of microbubble (MB)-assisted ultrasound (also known as sonoporation or sonopermeabilization) has gained momentum. The technique aims at delivering therapeutic molecules including nucleic acids, anti-cancer drugs, peptides and antibodies *in vitro* and *in vivo* [92,93]. It has been known for some time that ultrasound disturb the dynamic balance of the intracellular ROS, suggesting that the physical damage from sonoporation causes an increase in intracellular calcium which induces a rise of ROS in mitochondria. Recently, the exogenous molecular uptake and the concentration of intracellular ROS was quantified at the cellular level to evaluate the extent of the cell response to sonoporation [94]. The results suggest that that intracellular ROS generation in reversible sonoporated cells can impact their long-term fate.

Recently we have shown, using combined MD simulations and in vitro experiments, that under certain sonoporation conditions, ROS can form inside the microbubbles [57]. Experimental data confirmed theoretical predictions that MBs favor indeed spontaneous formation of a host of free radicals in which the hydroxyl radical (HO·) was the main species produced upon US exposure. The study has shown that these radicals could easily diffuse through the MB shell toward the surrounding aqueous phase and reach the nearby cells’ membranes.

An increase of both intracellular and extracellular ROS concentrations that might stem from the application of the sonoporation technique is shown to impact the permeability of the exposed cell. The exact mechanism is not fully described though one often assumes that cell membranes with extensive oxidation damage become leaky. Along the lines of previous simulations showing that HO· can penetrate deep in the hydrophobic core of lipids (POPC bilayers [31]) [57] to reach the peroxidation sites [95], namely their bis-allylic hydrogen atoms. Analyses using ROS scavengers and inhibitors suggested that the ROS produced under sonoporation conditions using MB-assisted ultrasound play a role in the permeabilization of cell plasma membranes and in the in vitro delivery of plasmid DNA [57]. The extent of permeability of cells to large molecules when subject to sonoporation is likely due (at least in part) to pores formed by secondary oxidation products, similar to those revealed in this study.

Evidently lipid oxidation is also implicated in a variety of other complex biological processes. Programmed cell death is a process that leads a living cell into controllable death that in contrast to necrosis, does not involve leakage of organelles neither induces proinflammatory response [96]. In recent year, many other forms of cell death aside of apoptosis were discovered, *e.g.* necroptosis or ferroptosis. The latter relates to an iron dependent mechanism involing its accumulation [97]. Ferroptosis is characterized by intensive ROS generation, lipid oxidation, NAPDH oxidation and GSH depletion [98,99]. Studies support a general model of cell disruption in necroptosis through the formation of small pores of a few nanometers radius in the plasma membrane [97],[100]. It was reported that, cells can be protected from ferroptosis by “pore blockage” using polyethylene glycol (PEG 1450 and 3350) [97], but not sucrose or raffinose [97] suggesting that the pores are larger than 1 nm and smaller than 2.4 nm in diameter. Pedrera *et al.* reported that ferroptosis progression involves production of peroxidized lipids prior to a sustained increase of cytosolic Ca^2+^ and final plasma membrane breakdown [101]. As high lipid peroxidationn is the most dramatic effect detected in ferroptosis, one could infer, consistently with the present work, that the pores detected are formed by local concentraitons of secondary lipid oxidation. Such pores’ formation in the cell plasma membranes can cause a significant drop in the transmembrane potential due to influx of calcium, sodium, chloride ions and outflux of potassium ions. Such changes in the ionic balance, might cause a depolarization of the cell and the activation of Voltage-Gated Calcium Channels (VGCC), and therefore further influx of calcium.

### 5. Conclusions

In summary, we investigated in this study whether the secondary lipid oxidation products could contribute to the long-lived permeability and conductance to the electropermeabilized cell lipid membranes, which persists after the application of the electric pulse. We performed a large number of simulations that show that large and long lifetime pores form due to the presence of a small patch of oxidized lipids. We calculated the permeability of such pores and that of bilayer models mimicking the topology of cell membranes well after pore closure. Overall, our modeling results and data analyses indicate that the pore formation due to the presence of secondary oxidation aldehyde lipids can quantitatively match the lowest to the highest reported experimental changes in the cells’ conductivity and permeation. Along with our previous report [35] this study therefore represents the first molecular level quantitative analyses of the permeation of cell membranes subject to oxidative damage confronted with experimental measurements.

We investigated the influence of membranes’ composition on the lifetime of the pores by studying mimics of lipid “gel-like domains” containing cholesterol concentrations and have shown that indeed the lipid dynamics in these cases slows down the pores’ annihilation. Real cell membranes are however crowded, with ~ 50% of the cell’s surface composed of proteins [102]. Furthermore, proteins show strong negative influence on lipid membrane’s fluidity [103,104], [105]. We can therefore safely conclude based on our investigations that the presence of secondary oxidation lipid patches resulting from direct oxidation could form pores with lifetimes far longer than herein assessed from simulations (or for instance from biophysical experiments on vesicles) where proteins are absent. Since starved cells don’t reseal completely [106], it is possible that some pores would remain formed as long as the cell lipid machinery does not take over to repair the damage.

As we have addressed in the discussion section, the relevance of these results is not restricted to electroporation-based treatments. Damage of the same topology and characteristics might be encountered whenever cells are exposed to extensive intracellular or extracellular ROS.

## Acknowledgments

The authors acknowledge the Centre de Calcul Régional *ROMEO (https://romeo.univ-reims.fr/)* for providing substantial computational resources.The authors thank Paulina Rozborska and Kinga Woźniak for the preliminary preparation of the patch systems and Lea Rems for assistance in the analyses of the free energy calculations.

## Appendices

Supporting Information

